# Comparison of the 3D organization of sperm and fibroblast genomes using the Hi-C approach

**DOI:** 10.1101/006247

**Authors:** Nariman Battulin, Veniamin S. Fishman, Alexander M. Mazur, Mikhail Pomaznoy, Anna A. Khabarova, Dmitry A. Afonnikov, Egor B. Prokhortchouk, Oleg L. Serov

## Abstract

The 3D organization of the genome is tightly connected to its biological function. The Hi-C approach was recently introduced as a method that can be used to identify higher-order chromatin interactions genome-wide. The aim of this study was to determine genome-wide chromatin interaction frequencies using the Hi-C approach in mouse sperm cells and embryonic fibroblasts. The obtained results demonstrated that the 3D genome organizations of sperm and fibroblast cells show a high degree of similarity both with each other and with the previously described mouse embryonic stem (ES) cells. Both A- and B-compartments and topologically associated domains (TADs) are present in spermatozoa and fibroblasts. Nevertheless, sperm cells and fibroblasts exhibited statistically significant differences between each other in the contact probabilities of defined loci. Tight packaging of the sperm genome resulted in an enrichment of long-range contacts compared with the fibroblasts. However, only 30% of the differences in the number of contacts are based on differences in the densities of their genome packages; the main source of the differences is the gain or loss of contacts that are specific for defined genome regions. An analysis of interchromosomal contacts in both cell types demonstrated that the large chromosomes showed a tendency to interact with each other more than with the small chromosomes and vice versa. We found that the dependence of the contact probability P(s) on genomic distance for sperm is in a good agreement with the fractal globular folding of chromatin. The similarity of the spatial DNA organization in sperm and somatic cell genomes suggests the stability of the 3D structure of genomes through generations.

## Introduction

For a long time, the study of chromosome architectures was based on fluorescence-based microscopy (Lanctôt et al. 2007; Joffe et al. 2010; Markaki et al. 2010). The approach allowed researchers to establish that individual chromosomes are localized in distinct spaces designated as chromosome territories (Cremer and Cremer 2001). Moreover, chromosome territories in nuclei are localized in a non-random manner with respect to the nuclear periphery (Cremer and Cremer 2001) and are able to interact and form gene clusters that loop out of their chromosome territory (Sproul et al. 2005). The development of a technique based on chromosome conformation capture (3C) (Dekker et al. 2002) and related methods (4C, 5C and Hi-C) (Zhao et al. 2006; Simonis et al. 2006; Dostie et al. 2006; Lieberman-Aiden et al. 2009) significantly extended the possibility of studying the three-dimensional (3D) genome architecture. The Hi-C technology, as a genome-wide approach, allows the determination of the contact frequency between any pair of loci within 10-100 nm at the moment of nuclei fixation (Dekker et al. 2002; 2013). Thus, Hi-C provides “a true all-by-all genome-wide interaction map” (Dekker et al. 2013b) based on the quantitative estimation of proximity-ligation events for millions of loci in the genome. Importantly, the Hi-C interaction frequencies are well correlated with the mean spatial distance separating loci, as measured using independent methods such as FISH (Eskeland et al. 2010; Kalhor et al. 2012), indicating that the Hi-C data can accurately reproduce the expected distance.

Genome-wide Hi-C mapping has revealed that inter- and intrachromosomal interactions are represented by two compartments, A and B, which have a mean size of approximately 5 Mb each (Lieberman-Aiden et al. 2009; Zhang et al. 2012; Imakaev et al. 2012). Loci of the A compartments interact preferentially with loci of other A compartments, while the B compartments often are in contact with other B compartments. Additionally, the genome-wide Hi-C mapping, in combination with a hidden Markov model, revealed that human and mouse chromosomes are composed of approximately 2,200 topologically associated domains (TADs) that have a median size of 880 kb and cover over 90% of the genome (Dixon et al. 2012). The same conclusion was simultaneously made based on the 5C analysis of the mouse X-chromosome inactivation center (Nora et al. 2012). It is important to note that the topological domains are stable across different cells (mouse embryonic stem (ES) cells and mouse cortex or human ES cells and human IMR90 fibroblasts) and highly conserved across species (human and mouse), “indicating that topological domains are an inherent property of mammalian genome” (Dixon et al. 2012).

In mammals, chromatin organization in mature sperm cells is unique among cell types. The genome of sperm cells is packaged in a highly condensed configuration. This packaging enables more than a ten-fold decrease in nucleus size in spermatozoa relative to the somatic interphase nucleus. This extraordinary compactness results from the replacement of histones with protamines. Protamines coil sperm DNA into toroids that form an almost crystalline structure. Only 1-15% of mammalian chromatin is bound to histones rather than protamines. Additionally, sperm cells have a haploid, transcriptionally inactive set of chromosomes (Johnson et al. 2011; Mudrak et al. 2011; Johnson et al. 2011). It is unknown how all of the aforementioned features affect the 3D organization of sperm genome.

The aim of this study is to compare the three-dimensional genome architectures of sperm cells and fibroblasts, as somatic cells, using the Hi-C approach. The obtained results demonstrate that “genome-wide interaction maps” of mouse sperm and fibroblast genomes show a high degree of similarity both to each other and to the previously described Hi-C organization of mouse ES cells. Nevertheless, there are statistically significant differences in the spatial contacts of some regions, e.g., in chromosomes 2, 5, 12, 13, and 19.

## Results

We created Hi-C libraries from mouse fibroblasts and mature sperm cells using the TCC-protocol developed by Kalhor and colleagues (Kalhor et al. 2012). The TCC method allows one to significantly reduce the noise obtained using the Hi-C approach, particularly the noise from interchromosomal interactions. We performed the massive parallel sequencing of the Hi-C libraries at a depth of 150 and 400 million read pairs for fibroblasts and sperm, respectively, and filtered the data so that the reads could be uniquely aligned to the mouse genome reference sequence.

Fig. 1 presents genome-wide and chromosome 19 Hi-C maps for sperm cells and fibroblasts (binned at 1 Mb resolution) as heatmaps, where the intensity indicates the contact frequency. Both interaction maps display visibly similar plaid patterns of the regional enrichment or depletion of long-range interactions. Individual chromosomes visually rise above both heatmaps due to the enrichment of contacts.

**Fig. 1.**
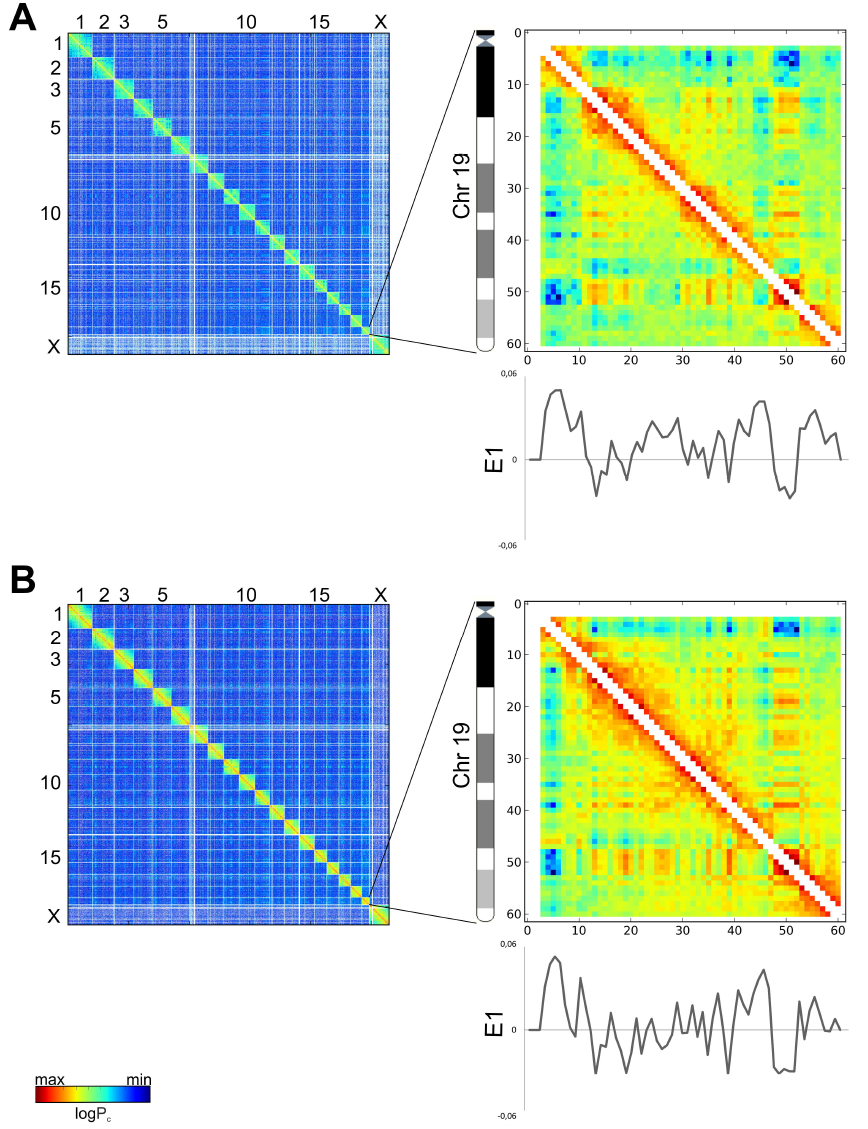
Relative Hi-C contact probability maps for whole-genome (on the left) and chromosome 19 (on the right) at a 1 Mb resolution for sperm cells (A) and fibroblasts (B). The two-dimensional iteratively corrected heatmaps demonstrate the characteristic plaid patterns for both sperm cells and fibroblasts. The color of each dot represents the log of the interaction probability for the corresponding genome bins. The graphs under the heatmaps show E1 values for chromosome 19 in sperm cells and fibroblasts.

A previous study showed that contact heatmaps could be decomposed into a set of eigenvector tracks. The first eigenvector (E1) values correlate with different genome properties such as replication time, GC-content and histone marks (Imakaev et al. 2012). We performed eigenvector decomposition and compared the obtained E1 values for spermatozoa, fibroblasts and ES cells using the Hi-C data published by Dixon and colleagues (Dixon et al. 2012) (Figs. 1, 2). Again, one can note a high degree of similarity between fibroblasts and spermatozoa (Spearman r = 0.898), while the level of similarity between fibroblasts and ES cells was slightly lower (Spearman r = 0.826). Lieberman-Aiden et al. (2009) suggested that the genome could be divided into discrete A- and B-compartments that are characterized by positive and negative E1 values or into a continuous set of domains that are characterized by similar E1 values for regions inside a compartment (Imakaev et al. 2012). Independent research supported this viewpoint (Dekker et al. 2013a). Thus, the similarities of E1 values imply a similar distribution of A- and B-compartments in sperm cells and fibroblasts, emphasizing conservatism in genome organization (Figs. 1, 2).

**Fig. 2.**
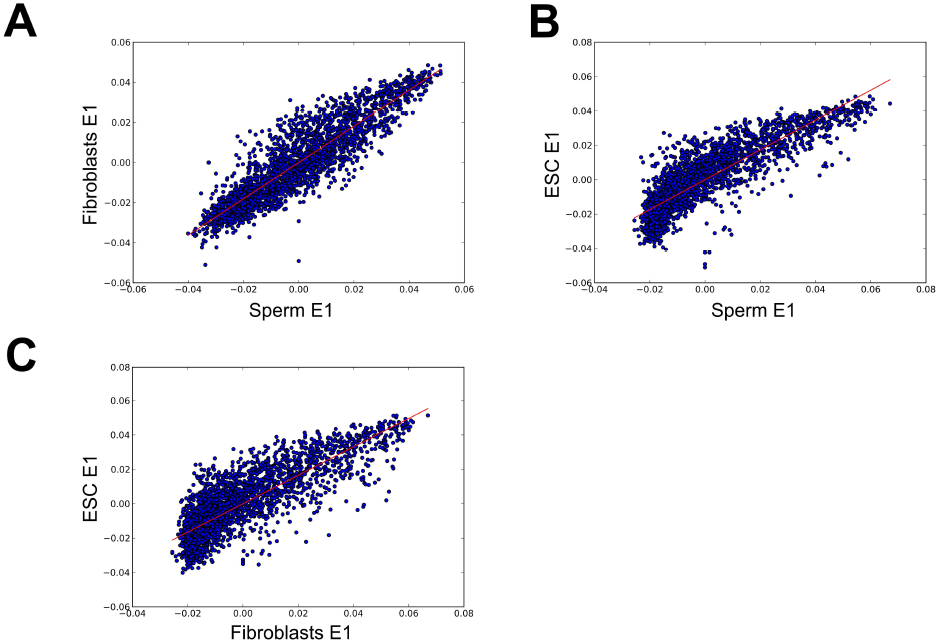
Comparison of the E1 values for both sperms cells and fibroblasts. Scatter plots of eigenvectors. The E1 values are highly similar in fibroblasts and sperm cells. The X- and Y-axis indicate the E1 values from fibroblasts and ES cells or fibroblasts and sperm. The line represents the linear trend for the obtained values.

In addition to the presence of A- and B-compartments, we identified TADs in sperm cells and fibroblasts (Fig. 3). We found 2590 domains in fibroblasts (with an average size of 928 kb and a median of 680 kb) and 1856 domains in sperm cells (with an average size of 1 226 kb and a median of 1000 kb).

**Fig. 3.**
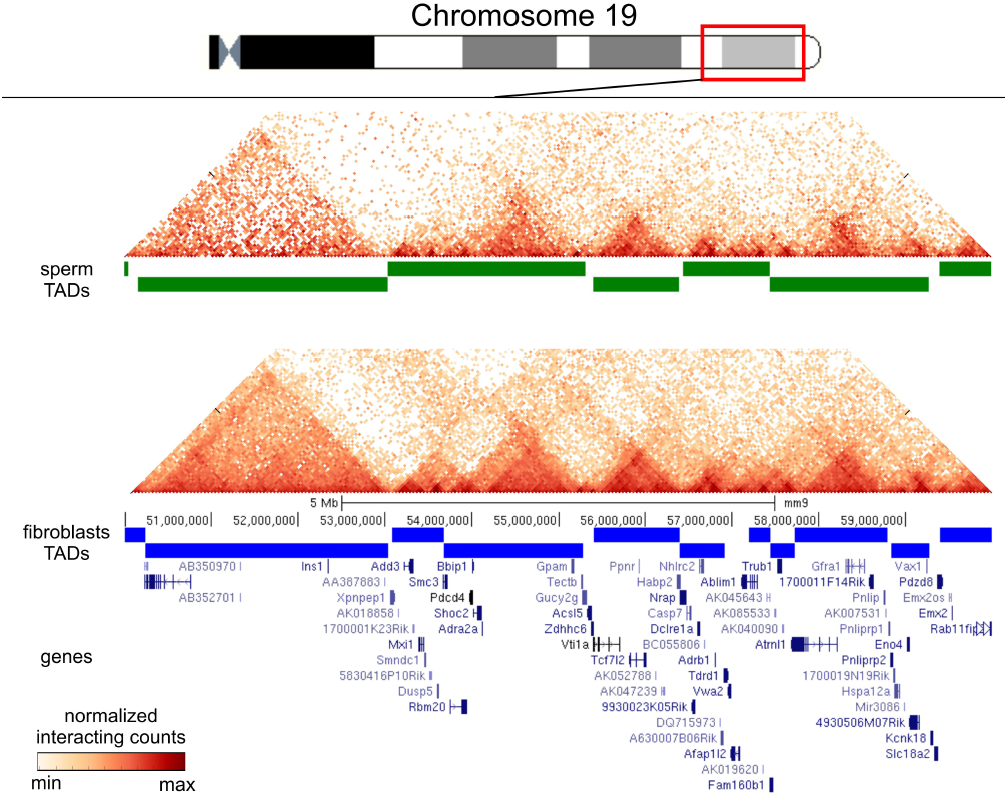
TADs are present in fibroblasts and sperm cells. The TAD signal is shown as a green line (for sperm cells) or a blue line (for fibroblasts) for a region on chromosome 19. The fragments of the heatmaps for sperm cells and fibroblasts (binned at a resolution of 40 kb) display the enrichment of contacts inside the TAD domains. The TAD signal shows visible similarity between sperm cells and fibroblasts.

The data above indicate that the general similarities of long-range interactions in sperm cell and fibroblast genomes do not exclude dissimilarities in any defined regions. Different analyses, such as the calculation of correlation coefficients, Euclid distance and comparisons of eigenvector values, have advantages and limitations and could even be connected to different biological properties; therefore, we decided to use all of these approaches together. We first compared the aforementioned characteristics (Pearson and Spearman correlation, Euclid distance and eigenvector values) for individual bins (Fig. 4, A). We observed a slight decrease in the Pearson and Spearman correlation coefficients for regions in the middle of chromosome compared with regions near the chromosome end. This difference was due to the statistical insignificance of rare long-range interactions captured for individual bins. We enhanced our comparison method to account for these biases using the self-correlating dataset of ES cells (see methods for details). Using this method, we selected, in sets of 100, the most different bins for each parameter separately, and we then focused on the bins that were present in all sets. We identified eight bins that greatly differ between fibroblasts and sperm cells (Table 1). The observed number of bins was more than 20 times higher than predicted (~0.1) for a random selection of bins. The identified regions that are dissimilar between sperm cells and fibroblasts are located on chromosomes 2, 5, 12, 13, and 19 (Table 1). Moreover, though large chromosomes have more bins and, therefore, a higher chance to be presented in the set of randomly selected bins, the smallest chromosome, 19, contains 3 of the 8 detected loci, indicating a non-random distribution of obtained regions.

**Fig. 4.**
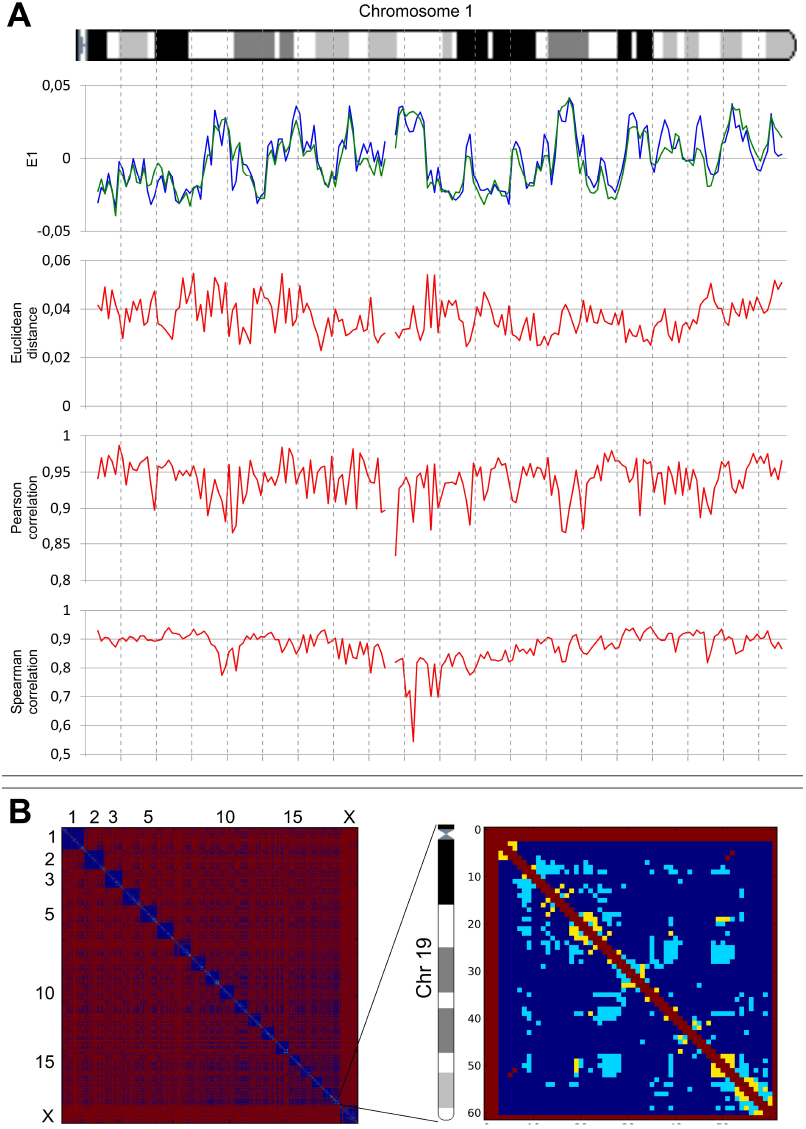
Identification of regions distinguishing sperm cells and fibroblasts. A. E1 values, Euclid distance and Pearson and Spearman correlation coefficients for chromosome 1 of both sperm cells and fibroblasts. All graphs indicate high similarities between sperm cells and fibroblasts (i.e., similar E1 values, small Euclid distance and high correlation coefficients). However, some regions display less similarity than others. B. The 2D heatmaps for the whole genome and for chromosome 19 (binned at a 1 Mb resolution) indicate the significance of the differences in the contact probability between fibroblasts and sperm cells. Each dot represents a single contact. Regions in red dots are not mappable, those in yellow dots are significantly different, those in cerulean are significantly different with a difference of more than 2 times, and those in blue are contacts where no significant difference was found.

**Table 1.** Localization of eight regions on a genome map, as identified by overlapping different datasets of highly dissimilar regions between sperm cells and fibroblasts

| Chromosome<br>number | Nucleotide start on mouse genome<br>map | Nucleotide end on<br>mouse genome map |
| --- | --- | --- |
| chr2 | 169000000 | 170000000 |
| chr5 | 119000000 | 120000000 |
| chr5 | 142000000 | 143000000 |
| chr12 | 35000000 | 36000000 |
| chr13 | 57000000 | 58000000 |
| chr19 | 54000000 | 55000000 |
| chr19 | 8000000 | 9000000 |
| chr19 | 9000000 | 10000000 |

We used another approach based on the identification of the individual contacts distinguishing sperm cells and fibroblasts to compare these cells. We considered all contacts supported by more than one read as mappable. Approximately 900 000 (~9%) of a potential ~7x10^−7^ (at a resolution of 1 Mb) contacts were mappable in sperm and fibroblast genomes. In addition, we found that of the 845 682 mappable interactions for both sperm cells and fibroblasts, 7 228 (0.85%) have significantly different (q-value < 0.05) contact probability. Moreover, of these 7 228 interactions, the probabilities of 5 930 contacts showed more than a two-fold difference (Fig. 4 B). Interestingly, the above-mentioned loci of chromosome 19 show a high amount of significantly different interactions with other regions in the genome (Fig. 4 B).

The dependence of the contact probability of genome loci on the distance between these loci P(s) is informative for the understanding of the DNA state (Naumova et al. 2013; Imakaev et al. 2012; Lieberman-Aiden et al. 2009). We examined the P(s) dependence in sperm cells and fibroblasts. For both cell types, we observed a strong decrease in contact probability with an increase in the distance between loci, i.e., P(s)~s^1,07^ for spermatozoa and P(s)~s^1,27^ for fibroblasts (Fig. 5, A). Interestingly, the contact probability for fibroblasts was higher, in the range of 10^4−^10^−6^ bp. This increase in P(s) values was compensated by a lower contact probability in a diapason of long-range interactions at 10^−7^-10^−8^ bp. These data suggest that sperm cells have more long-range contacts than do fibroblasts. A detailed analysis showed that the probabilities of contacts in fibroblasts were less than those in sperm cells, when counting regions separated by less than 40 Mb; for loci separated by 50-150 Mb, fibroblasts display more than two times higher probabilities of contacts compared with sperm cells (Fig. 5, B). Overall, these data indicate that the genome of spermatozoa is packed more compactly, such that more distant loci are brought together and have a high probability of contact with each other.

**Fig. 5.**
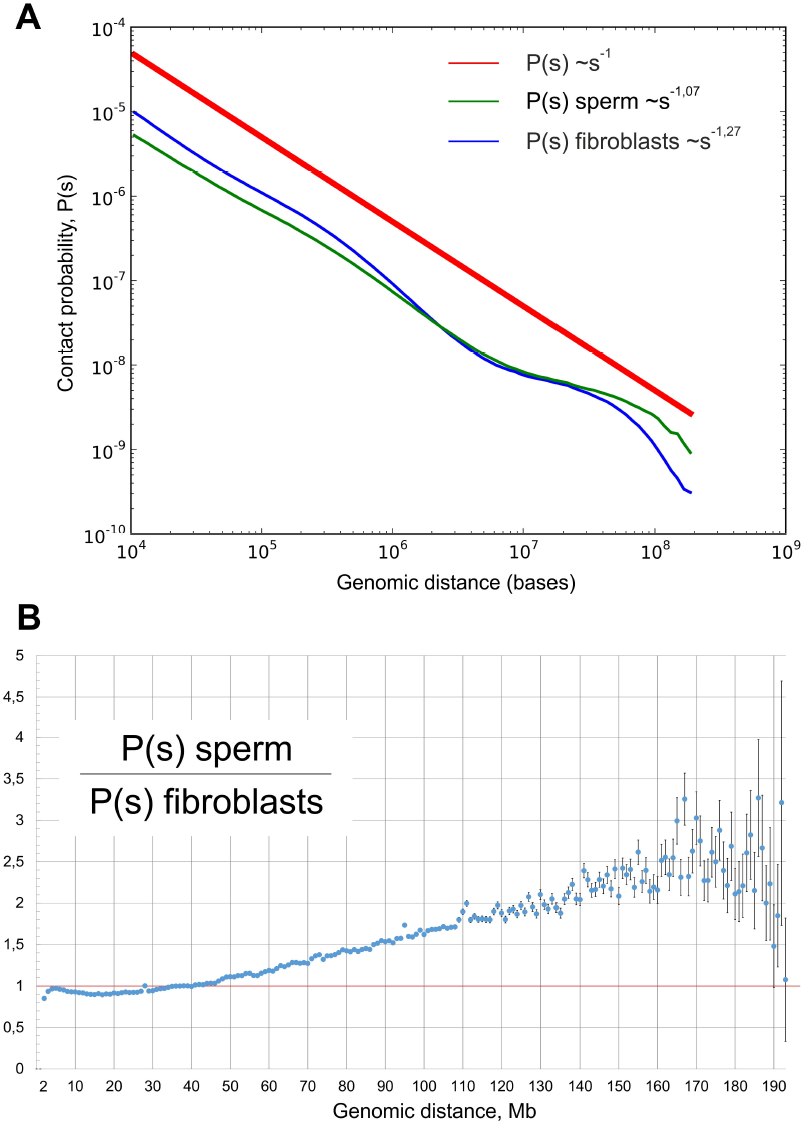
The genome of sperm cells is packed more tightly than that of fibroblasts. A. The dependence of the contact probability on the genomic distance P(s) averaged over all chromosomes, compared with P~1/s. The blue line indicates fibroblasts (P~s^−127^), and the green line indicates sperm cells (P~s^−107^). B. The ratio between sperm cells’ and fibroblasts’ contact probabilities at different genomic distances. The X-axis indicates genomic distance, and the Y-axis indicates the ratio of contact probabilities. The black lines show a 1:1 ratio.

Consistent with an increased amount of long-range interactions, sperm cells showed higher intrachromosomal to interchromosomal contacts ratio than did fibroblasts (Fig. 6 A). We observed 25-40 times more intrachromosomal contacts than interchromosomal ones in fibroblasts, whereas sperm cells showed only a difference of 12-20 times more intrachromosomal contacts.

**Fig. 6.**
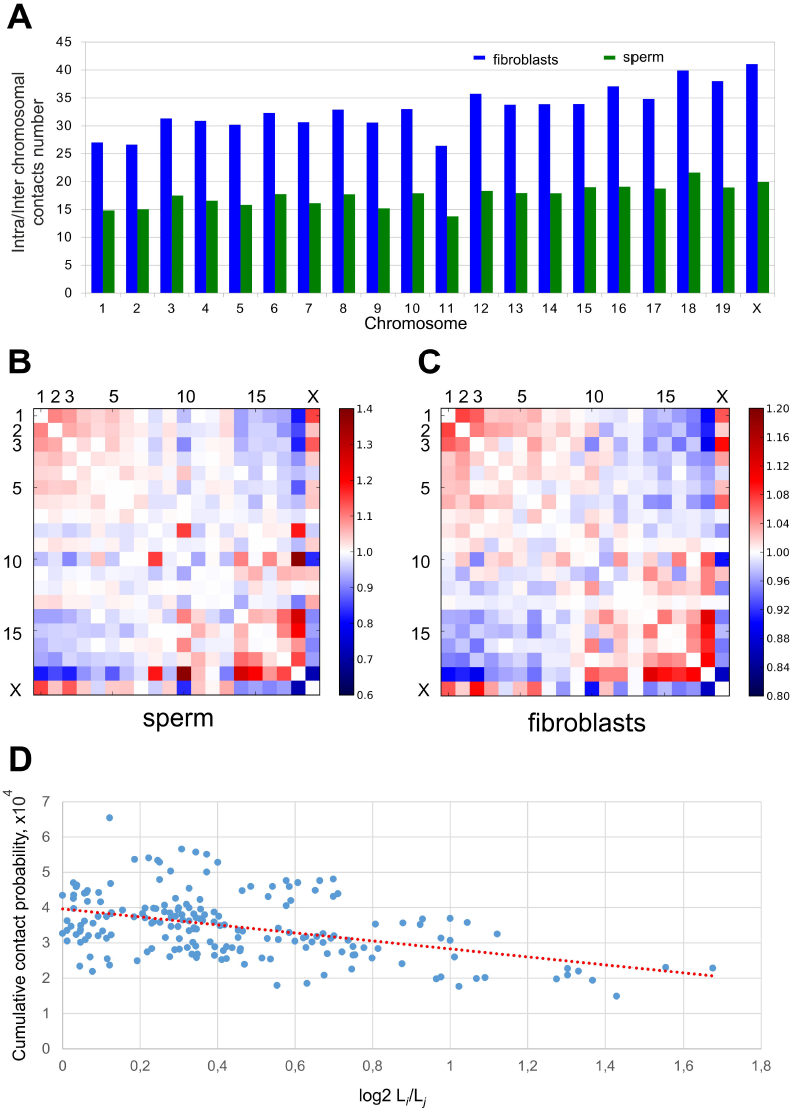
Analysis of intrachromosomal contacts in sperm cells and fibroblasts. A. The ratio between intra- and interchromosomal contact numbers for sperm cells (green) and fibroblasts (red). B, C. The 2D heatmaps show the observed number of interactions between any pair of chromosomes divided by the expected number of interactions between those chromosomes for sperm cells (B) and fibroblasts (C). The color of each dot represents the enrichment (red) or depletion (blue) of contacts compared with the expected values. D. The observed number of interactions between any pair of chromosomes plotted against the difference in the lengths of those chromosomes. The dotted lined represents the linear trend for obtained values.

In the generated heatmaps, individual chromosomes showed up as compact contact-enriched clusters (Fig. 1). Indeed, more than 90% of the interactions fell into interchromosomal contacts in both sperm and fibroblast cells. Statistical analyses of rare interchromosomal contacts revealed that chromosomes are distributed non-randomly in sperm and fibroblast nuclei (Fig. 6 B). The whole chromosome interaction patterns show that the large chromosomes (for instance, chromosomes 1-8 and X) are more likely to interact with one another and not with the small chromosomes (chromosomes 14-19), while the shorter chromosomes show a tendency to establish contact with each other (Fig. 6 B). A similar pattern of interactions between chromosomes was identified in fibroblasts (Fig. 6 C). This observation was further confirmed by an opposite correlation between probabilities of contacts between chromosomes and differences in their lengths (Pearson r=-0.44) (Fig 6, D). These data are in agreement with the previously published Hi-C results (Zhang et al. 2012).

The differences between the 3D organizations of sperm cells and fibroblasts can potentially originate from two independent sources. First, the denser packaging of sperm genome compared with the fibroblast genome is due to a decrease in the nucleus size and a denser packaging of DNA with protamines compared with histones. Second, local rearrangements of 3D genome structures are due to the loss or gain of functional connections between different loci. To estimate the role of the first reason (i.e., the denser packaging of the sperm genome), we developed a model process referred to as the “compression” of the fibroblast genome to a sperm cell’s parameters. Our model does not change the distribution of contact probabilities for regions separated by the same genomic distance but instead brings all loci closer to each other. Thus, we normalized the number of fibroblast contacts between loci separated by a given genomic distance to achieve the same P(s) distribution for fibroblasts and sperm cells, but we maintained the contact ratios of loci separated by the same genomic distance. We compared the obtained post “compression” fibroblasts (C_Sp_-fibroblasts) with the sperm cells and found that the number of contacts that have different probabilities between sperm cells and fibroblasts decreased from 7 228 to 5 106 after “compression”. Additionally, the number of contacts, with more than a two old difference in contact probabilities, decreased from 5 930 to 4 278. As a control, we performed the “compression” of fibroblasts to ES cell parameters, thereby obtaining C_ESC_-fibroblasts. The number of differences in contact probabilities between C_ESC_-fibroblasts and sperm cells increased up to 11 524, with more than 10 272 contacts showing at least a two-fold difference. However, the C_ESC_-fibroblasts were more similar to ES cells than original fibroblasts were to ES cells, indicating that “compression” decreases differences only when performed in a cell-specific manner. Our data imply that approximately 30% of the differences in contact probabilities between sperm cells and fibroblasts might originate from differences in the densities of their genome packaging; however, the main source of differences is the gain or loss of contacts that are specific to defined genome regions.

## Discussion

The obtained data are the first description of 3D organization in mouse motile sperm cells and fibroblasts obtained using Hi-C technology. Though spermatozoa and fibroblasts are extremely different in a number of aspects, the spatial organization of DNA in these cells is similar. Moreover, two types of previously identified domains, i.e., A- and B-compartments (Lieberman-Aiden et al. 2009; Imakaev et al. 2012) and TADs (Dixon et al. 2012), were present in both sperm and fibroblast genomes.

It is still unknown whether the presence of spatial domains in cells is an indirect result of DNA packaging in nucleosomes and the transcription process or whether there are special mechanisms involved in the formation and maintenance of spatial domains. In sperm cells, DNA packaging is influenced at a very basic level by the replacement of histones with different proteins, i.e., protamines; the transcription process is also completely abolished (Mudrak et al. 2011). However, the high-order chromatin structure of the cell remains stable. This finding suggests the presence of special mechanisms involved in the establishment and maintenance of these structures and highlights an important role for spatial domains in cell function.

Despite the remarkable similarity of 3D genome organization between sperm cells and fibroblasts, we aimed to find regions that distinguish these cell types. We used three independent methods (Pearson correlation, Euclidian distance and eigenvector comparison) to compare the 3D organization of the genomes of sperm cells and fibroblasts. Some of these methods (Pearson correlation, and eigenvector comparison) have been used previously (Imakaev et al. 2012; Lieberman-Aiden et al. 2009); we introduced Euclidian distance as a method for comparing individual genomic regions. Though the overlap between the sets of genomic regions obtained using different methods was more than 100 times larger than expected for randomly selected regions, it was still far from 100%. One explanation for this result can be a difference in the sensitivities of the methods used to systematic biases of the Hi-C experiment. Another, more intriguing explanation is that different the mathematical methods used to compare individual regions reflect different biological properties of these regions. Additional studies are required to develop a standardized approach for the comparison of several Hi-C datasets.

To explain the differences observed in sperm cells and fibroblasts, we developed a genome compression model. This model differs from those described by Mirny and colleagues (Naumova et al. 2013; Mirny 2011). While the approaches developed by Mirny and colleagues are based on the modeling of polymer region at a physical level (Naumova et al. 2013; Mirny 2011), our model is based on the mathematical operations obtained in experimental Hi-C matrices. Our model shows that genome compression can explain approximately 30% of the differences between sperm cells and fibroblasts. The further development of such a model might allow understanding the changes in 3D-structure of chromatin from a new point of view.

The differences described above, i.e., in the long-range contacts in sperm cells and fibroblasts, are most likely related to the extreme compactness of the sperm genome. In fact, the DNA within the sperm nucleus is packed in a volume that is approximately 5% of the volume in somatic cells (Mudrak et al. 2011). Here, it is pertinent to note that the compactness of the sperm genome is comparable to that of metaphase chromosomes. Recently, Naumova et al. (Naumova et al. 2013) reported a homogenous folding state that is locus-independent and common to all chromosomes at their metaphase status in examined cell types. Keeping in mind the similarity of the 3D organization of sperm cells and fibroblasts, one could suggest that the exceptional compactness of the sperm genome is not sufficient in itself to change hierarchical models of chromatin structure (Imakaev et al. 2012; Dixon et al. 2012; Dekker et al. 2013a).

Consistent with the above reasoning, we found the P(s) distribution in sperm cells to be in agreement with a fractal model of genome organization. The P(s) distributions in spermatozoa and fibroblasts were strictly different from those found in mitotic chromosomes (Naumova et al. 2013), emphasizing differences in the mechanisms of genome compression during mitosis and sperm maturation.

In summary, the remarkable similarities in the 3D genome organization of spermatozoa and fibroblasts show the role of male gametes as carriers of the 3D genome organization through generations.

## Methods

### Preparation of motile sperm cells and mouse embryonic fibroblasts

Mature mouse spermatozoa were obtained from the epididymis of C57BL mice using the swim-up assay (Brykczynska et al. 2010). Briefly, *cauda epdidymis* were dissected into pieces and placed into sperm motility medium (135 mM NaCl, 5 mM KCl, 1 mM MgSO_4_, 2 mM CaCl_2_, 30 mM Hepes, pH 7.4; freshly supplemented with 10 mM lactate acid, 1 mM sodium pyruvate, 20 mg/ml bovine serum albumin (BSA), 25 mM NaHCO_3_) for 1 hr at 37°C. To avoid contamination by somatic cells, only the top fractions containing motile sperm were collected. The cell suspension was centrifuged, and the cell pellets were resuspended in serum free Dulbecco MEM medium and processed for Hi-C library generation as described below.

Mouse embryonic fibroblasts were obtained from 13-day-old embryos from C57BL mice and cultured in standard conditions, as described previously (Kruglova et al. 2008).

### Generation of Hi-C libraries

Hi-C libraries were produced using tethered conformation capture protocol (Kalhor et al. 2012), but with some minor modifications. Briefly, 50 million sperm cells were resuspended in 45 ml serum free Dulbecco MEM medium, and 37% formaldehyde was added to obtain a final concentration of 1% for cross-link chromatin. Mouse fibroblasts were fixed while they were attached to the culturing surface in 1% final concentration of formaldehyde in the serum free Dulbecco MEM medium. Cells were incubated at room temperature for 10 min; the formaldehyde was then quenched by adding glycine to obtain a final concentration of 0.125 M, and the mixture was incubated at room temperature for 5 min and subsequently on ice for 15 min. Mouse fibroblasts were scraped from the culture plate using disposable cell scrapers and aliquoted for 25 million cells. Sperm cells were harvested by centrifugation. After crosslinking, the sperm and fibroblast samples were processed identically. Fixed cells were lysed using a Dounce homogenizer in the presence of cold lysis buffer (10 mM HEPES, pH 8.0, 10 mM NaCl, 0.2% IGEPAL CA-630, and 1X protease inhibitor solution).

The chromatin was solubilized with dilute sodium dodecyl sulfate (SDS) and incubation at 65 °C for 10 min. The chromatin was biotin labeled chemically by EZ-Link-Iodoacetyl-PEG2-biotin (Pierce Protein Research Products). DNA in the cross-linked protein complexes were digested with *HindIII* endonuclease. Biotinylated digested chromatin was immobilized on MyOne Streptavidin T1 beads (Invitrogen). The 5’ overhang was filled in by the Klenow fragment of DNA polymerase I using equimolar amounts of all deoxyribonucleotides, with the substitution of biotin-14-dCTP for dCTP. The immobilized blunt-ended DNA fragments were then ligated while they were tethered to the surface of the beads. The chromatin complexes containing the biotin-labeled ligation products were degraded by incubation with Proteinase K at 65 °C. DNA was purified by phenol-chloroform extraction. The biotinylated nucleotide was removed from non-ligated DNA ends using T4 DNA polymerase, as previously described (Belton et al. 2012). The DNA was sheared and size-selected; the fragments that included a ligation junction were then isolated on streptavidin-coated magnetic beads and prepared for paired-end sequencing. The libraries were sequenced on an Illumina Genome Analyzer IIx (GA IIx) machine using the paired-end module and with 50 bp reads on each end.

### Generation of heatmaps

Sequencing reads were mapped to the mm9 mouse genome and filtered using the pipeline developed by Imakaev et al (Imakaev et al. 2012). Mirnlib version 0d30147f052f and hiclib version d28d8d985120 software was obtained from http://mirnylab.bitbucket.org/hiclib/. The public datasets SRR443883, SRR443884 and SRR443885 (Dixon et al. 2012) were processed similarly to obtain Hi-C data for mouse ES cells. Heatmap computation, iterative correction, eigenvector decomposition and P(s) calculation were performed using the hiclib software (Imakaev et al. 2012). For each interaction, we estimated an error as 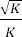, where K is a number of reads supporting the interaction. The average error for non-zero intrachromosomal interactions in sperm cells was 24% at the 1-megabase (MB) scale and reached 88% on a 0.1-MB scaled heatmap. Therefore, we used the 1 MB resolution for all subsequent calculations, except cases where the resolution is specifically indicated.

### Identification of regions different between sperm cells and fibroblasts

We calculated the Euclidean distance, Pearson correlation and eigenvector differences between individual bins of sperm and fibroblasts cells.

We calculated Euclidean distance between individual bins as

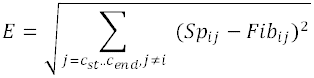

where E is the Euclidean distance between bin “i” at chromosome chr_i_, c_st_ is the first bin of chromosome chr_i_, c_end_ is the last bin of chromosome chr_i_, and Sp_ij_ and Fib_ij_ are the number of reads supporting contacts between bins “i” and “j” of sperm cells and fibroblasts. A greater Euclidean distance indicates a larger difference between bins.

The Pearson and Spearman correlation coefficients were calculated for each bin, accounting only for intrachromosomal contacts. The signal-to-noise ratio might vary in a Hi-C experiment, even if the data are iteratively corrected. The enrichment of any genomic region with interactions that have low signal-to-noise ratios might result in an underestimation of the correlation coefficients for these genomic regions. To handle this problem, we used the correlation of two random sub datasets (“reference” datasets) generated from the ES cells dataset as a marker for regional-dependent biases. We defined poor bins as those that satisfy the condition

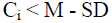

where C_i_ is the Pearson or Spearman correlation coefficient between bin “i” in the “reference” datasets, and M and SD are the median and standard deviation of the Pearson correlation coefficient for all bins in the “reference” datasets. We then excluded from the analysis all poor bins as well as 3 bins both upstream and downstream of each poor bin. Because the Spearman correlation coefficient is rank-based, it is more sensitive to small differences in the samples observed at low signal-to-noise ratios. We therefore used the Pearson correlation coefficient for the subsequent analysis.

The 1^st^ eigenvector (E1) values of the sperm and fibroblast cells were calculated using the hiclib software, as described previously (Imakaev et al. 2012). We considered each bin to be a dot, with the X-coordinate equal to the appropriate E1 value of sperm cells and the Y-coordinate equal to the appropriate E1 value of fibroblasts. We computed a linear regression line for the obtained dots using the least-squares method. We than calculated a distance from each dot to the regression line and used this distance as a measure of the difference between the eigenvectors of the appropriate bins. A greater distance indicates a larger difference between bins.

We ranked all bins in the sperm cell and fibroblast datasets using 3 types of ranks (ranks generated during the calculations of Euclidean distance, Pearson correlation and eigenvectors difference). From the 2308 bins, we selected the 100 highest-ranked bins for each type of analysis and defined them as candidate bins. This process resulted in 3 sets of candidate bins. Some of the candidate bins might have had high ranks due to region-specific biases (e.g., described in the calculation of Pearson correlation coefficient). To exclude such regions, we calculated the Euclidean distance and the difference between the eigenvectors for two “reference” ES cell datasets (described above in the calculation of Pearson correlation coefficient), selected 100 highest-ranked bins for each type of analysis and excluded them from the candidate bins. Finally, we identified regions that were present in all 3 sets’ candidate bins and defined them as regions that differed between sperm cells and fibroblasts. The expected number of regions that differed between sperm cells and fibroblasts was calculated as

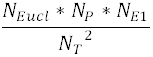

Where *N_Euc1_*, *N_P_*, *N_E1_* are the number of bins remaining after filtering candidate bins with ranks according to Euclidean distance *(N_Euc1_),* Pearson correlation *(N_P_)* and eigenvectors difference *(N_E1_)* and *N_T_* is the total number of bins (2308).

### Identification of differences in individual contact probabilities in sperm cells and fibroblasts

We used a uniform probability model to describe the contact frequencies observed in Hi-C experiments (Duan et al. 2010). Assuming that the probability of observing any particular interaction is uniform, the probability of contacts between bins “i” and “j” (*P^i,j^*) is

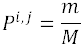

where m is the number of reads supporting the interaction (normalized in the iterative correction) and M is the total number of reads (normalized in the iterative correction). Note that when counting reads, we only considered contacts that were supported by more than one read (mappable contacts). We tested the null hypothesis *H_0_:* 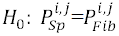, where 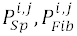 are the probabilities of contacts between bins “i” and “j” in sperm cells (P_Sp_) and fibroblasts (P_Fib_). Assuming normal approximation of binomial distribution, we calculated the p-value for the null hypothesis as

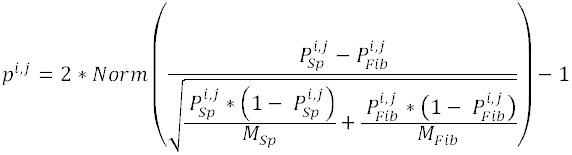

where *Norm* is the normal distribution. We than calculated q-values by multiplying each p-value by the total number of hypotheses tested.

### Modeling of fibroblast genome “compression”

To perform the fibroblast genome “compression”, we first calculated compression coefficients K_j_ as

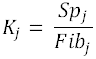

Where Sp_j_, Fib_j_ represent the sum of elements of diagonal j in iteratively corrected Hi-C matrix for sperm cells or fibroblasts, respectively. Errors for coefficient calculation were estimated in the same way as described above (in Generation of heatmaps). We then performed a correction by multiplying all contacts at the diagonal j of the fibroblast datasets by the appropriate coefficient compression coefficients. We did not apply a correction if the coefficient error was above 5%. Finally, we adjusted all contacts to achieve the same total sum of elements for both the resulting and original matrices.

The chromosome interaction patterns were calculated as described previously (Kalhor et al. 2012; Lieberman-Aiden et al. 2009). Briefly, the observed/expected contact frequencies for chromosomes “i” and “j” were calculated as

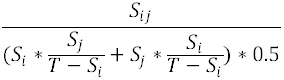

where S_i_ and S_j_ are the sum of interchromosomal contacts of chromosomes “i” and “j”, respectively, S_ij_ is the sum of contacts between chromosomes “i” and “j”, and T is the total sum of all interchromosomal contacts.

### Identification of TADs

To identify TADs, heatmaps were binned at 40 KB resolution, iteratively corrected and analyzed using a previously developed pipeline (Dixon et al. 2012).

## Data access

The sequencing results of Hi-C libraries of sperm cells and fibroblasts are available in the NCBI SRA archive under accession number SUB540202 (SRX553176 for sperm cells data and SRX554530 for fibroblasts data)

## Acknowledgments

The authors are indebted to Professors Nikolay Kolchanov and Nikolay Rubtsov from the Institute of Cytology & Genetics (Novosibirsk) for their valuable advice and comments at the initiation of this study. The study was partially supported by RFBR grant N<u>o</u> 14-04-31367 and and Skolkovo Center for Stem Cell Research.

